# Rewiring of liver diurnal transcriptome rhythms by triiodothyronine (T_3_) supplementation

**DOI:** 10.1101/2022.04.28.489909

**Authors:** Leonardo Vinícius Monteiro de Assis, Lisbeth Harder, José Thalles Lacerda, Rex Parsons, Meike Kaehler, Ingolf Cascorbi, Inga Nagel, Oliver Rawashdeh, Jens Mittag, Henrik Oster

**Affiliations:** Institute of Neurobiology, Center of Brain Behavior & Metabolism, University of Lübeck, Germany; Division of Molecular Neurobiology, Department of Medical Biochemistry and Biophysics, Karolinska Institutet, Stockholm, Sweden; Institute of Bioscience, Department of Physiology, University of São Paulo, Brazil; Australian Centre for Health Services Innovation and Centre for Healthcare Transformation, School of Public Health and Social Work, Faculty of Health, Queensland University of Technology, Kelvin Grove, Australia; Institute of Experimental and Clinical Pharmacology, University Hospital Schleswig- Holstein, Campus Kiel, Germany; School of Biomedical Sciences, Faculty of Medicine, University of Queensland, Brisbane, Australia; Center of Brain Behavior & Metabolism, Institute for Endocrinology and Diabetes – Molecular Endocrinology, University of Lübeck, Germany

**Keywords:** thyroid hormones, liver, hyperthyroidism, transcriptome, circadian clock

## Abstract

Diurnal (i.e., 24-hour) physiological rhythms depend on transcriptional programs controlled by a set of circadian clock genes/proteins. Systemic factors like humoral and neuronal signals, oscillations in body temperature, and food intake align physiological circadian rhythms with external time. Thyroid hormones (THs) are major regulators of circadian clock target processes such as energy metabolism, but little is known about how fluctuations in TH levels affect the circadian coordination of tissue physiology. In this study, a high triiodothyronine (T_3_) state was induced in mice by supplementing T_3_ in the drinking water, which affected body temperature, and oxygen consumption in a time-of-day dependent manner. 24-hour transcriptome profiling of liver tissue identified 37 robustly and time independently T_3_ associated transcripts as potential TH state markers in the liver. Such genes participated in xenobiotic transport, lipid and xenobiotic metabolism. We also identified 10 – 15 % of the liver transcriptome as rhythmic in control and T_3_ groups, but only 4 % of the liver transcriptome (1,033 genes) were rhythmic across both conditions – amongst these several core clock genes. In-depth rhythm analyses showed that most changes in transcript rhythms were related to mesor (50%), followed by amplitude (10%), and phase (10%). Gene set enrichment analysis revealed TH state dependent reorganization of metabolic processes such as lipid and glucose metabolism. At high T_3_ levels, we observed weakening or loss of rhythmicity for transcripts associated with glucose and fatty acid metabolism, suggesting increased hepatic energy turnover. In sum, we provide evidence that tonic changes in T_3_ levels restructure the diurnal liver metabolic transcriptome independent of local molecular circadian clocks.

## INTRODUCTION

Circadian clocks play an essential role in regulating systemic homeostasis by controlling, in a time-dependent manner, numerous biological processes that require alignment with rhythms in the environment (Gerhart-Hines and Lazar 2015; West and Bechtold 2015; de Assis and Oster 2021). At the molecular level, the clock machinery is comprised of several genes that are organized in interlocked transcriptional-translational feedback loops (TTFLs). The negative TTFL regulators, *Period* (*Per1-3*) and *Cryptochrome* (*Cry1-2*), are transcribed after activation by Circadian Locomotor Output Cycles Kaput (CLOCK) and Brain and Muscle ARNT-Like 1 (BMAL1 or ARNTL) in the subjective day. Towards the subjective night, PER and CRY proteins heterodimerize and, in the nucleus, inhibit BMAL1/CLOCK activity. This core TTFL is further stabilized by two accessory loops comprised by Nuclear Receptor Subfamily 1 Group D Member 1-2 (NR1D1-2, also known as REV-ERBα-β) and Nuclear Receptor Subfamily 1 Group F Member 1-3 (NR1F1-3, also known as ROR*α-γ*), and the *PAR-bZip* (proline and acidic amino acid-rich basic leucine zipper) transcription factor DBP (Albumin D-Site Binding Protein) (Takahashi 2017; Pilorz et al. 2020; de Assis and Oster 2021). Upon degradation of PER/CRY, towards the end of the night, BMAL1/CLOCK are disinhibited, and a new cycle starts.

How the molecular clocks in different tissues and downstream physiological rhythms are coordinated has been the subject of increasing scientific interest in recent years. Environmental light is detected by a non-visual retinal photoreceptive system that innervates the central circadian pacemaker, the suprachiasmatic nucleus (SCN) (Golombek and Rosenstein 2010; Hughes et al. 2016; Ksendzovsky et al. 2017; Foster et al. 2020). The SCN distributes temporal information to other brain regions and across all organs and tissues (Husse et al. 2015; de Assis and Oster 2021) through partially redundant pathways, including nervous stimuli, hormones, feeding-fasting, and body temperature cycles. Despite an ongoing discussion about the organization of systemic circadian coordination, all models share the need for robustly rhythmic systemic time cues (de Assis and Oster 2021).

The thyroid hormones (THs), triiodothyronine (T_3_) and thyroxine (T_4_), are major regulators of energy metabolism. In the liver, THs regulate cholesterol and carbohydrate metabolism, lipogenesis, and fatty acid ß-oxidation (Sinha et al. 2014; Ritter et al. 2020). While circadian regulation of the upstream thyroid regulator TSH (thyroid-stimulating hormone) has been described, T_3_ and T_4_ rhythms in the circulation show relatively modest amplitudes in mammals, probably due to their long half-life (Weeke and Laurberg 1980; Russell et al. 2008; Philippe and Dibner 2015). Interestingly, in hyperthyroid patients, non-rhythmic TSH secretion patterns are observed (Ikegami et al. 2019).

In this study, we investigated how a high T_3_ state in mice affects diurnal transcriptome organization in the liver. Our data show that tonic endocrine state changes rewire the liver transcriptome in a time-dependent manner independent of the liver molecular clock. Main targets of TH signaling are genes associated with lipid, glucose, and cholesterol metabolism.

## RESULTS

### Effects of high T_3_ on behavioral and metabolic diurnal rhythms

We used an experimental mouse model of hyperthyroidism by supplementing the drinking water with T_3_ (0.5 mg/L in 0.01 % BSA). Control animals (CON) were kept under the same conditions with 0.01 % BSA supplementation (Sjögren et al. 2007; Vujovic et al. 2015). TH state was validated by analyzing diurnal profiles of T_3_ and T_4_ levels in serum. Significant diurnal (i.e., 24-hour) rhythmicity was detected for T_3_ in CON with peak concentrations around the dark-to-light phase transition. T_3_ supplemented mice showed ca. 5-fold increased T_3_ levels compared to CON mice with no significant diurnal rhythm. T_4_ levels were non-rhythmic in all groups (Fig. 1A – B, Table S1). Compared to CON, overall T_4_ levels were reduced 2 to 3-fold in T_3_ supplemented animals (Fig. 1B). Resembling the human hyperthyroid condition, T_3_ mice showed increased average body temperature (Fig. S1A) as well as food and water intake compared to CON mice (Fig. S1B – C). Conversely, T_3_ mice showed higher body weight on the 3^rd^ week of experimentation (Fig. S1D), as previously shown (Johann et al. 2019).

**Figure 1:**
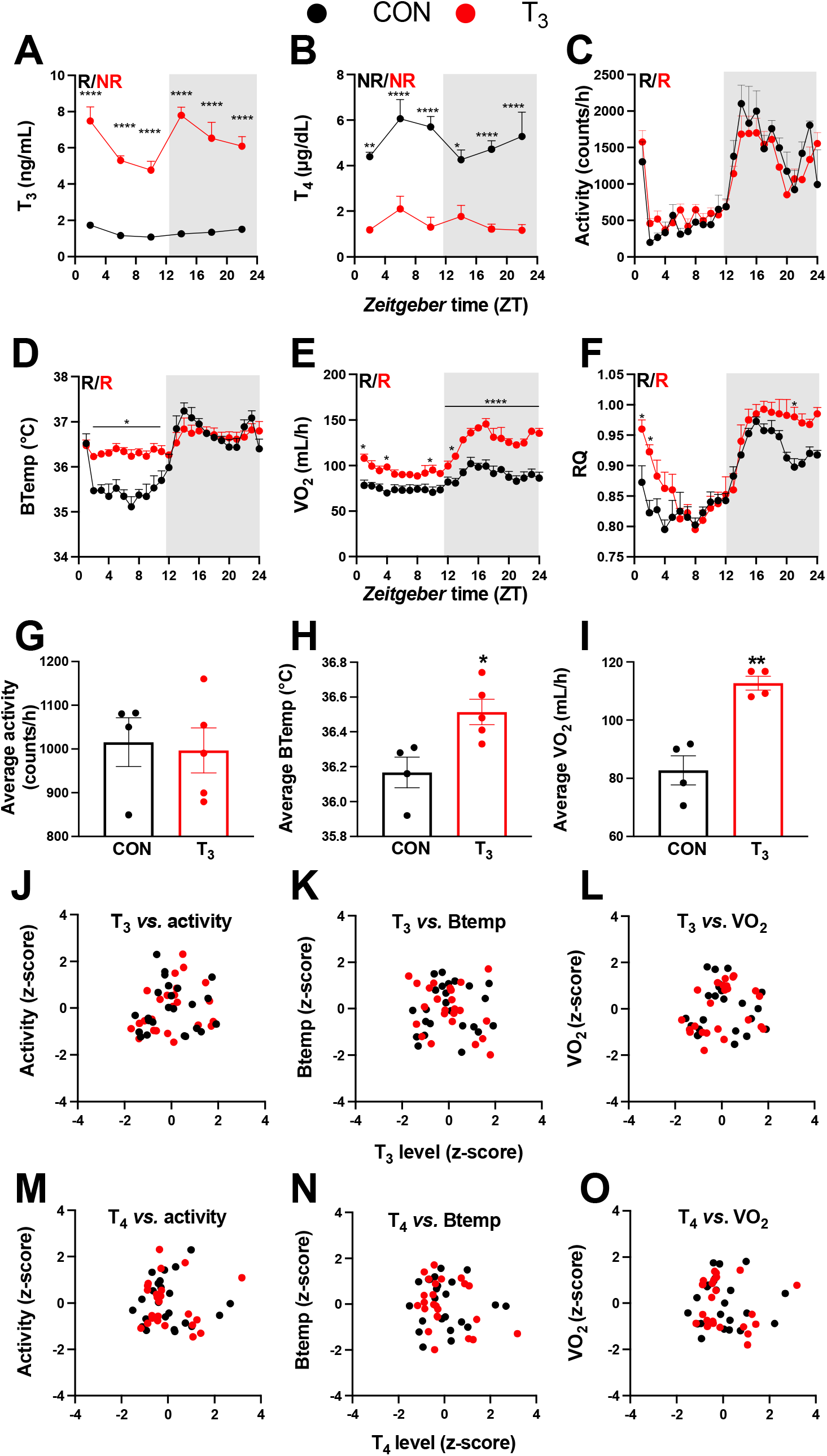
T_3_ treated mice show classic effects of high thyroid hormone levels compared to control mice (CON). A – F) Serum levels of T_3_ and T_4_, 24-hour profiles of locomotor activity, body temperature, O_2_ consumption and respiratory quotient are shown. Rhythm evaluation was performed by JTK cycle (p < 0.01, Table S1). Presence (R) or absence of circadian rhythm (NR) is depicted. G – I) Average levels of locomotor activity, temperature, and O_2_ consumption. J – O) Correlation between thyroid hormone levels and normalized levels of metabolic outputs are shown as z-scores (additional information is described in Table S2). In A and B, n = 4 – 6 animals per group and/or timepoint. In C and D, n = 4 and 5 for CON and T_3_ groups, respectively. In E and F, n = 4 for each group.

Metabolism-associated parameters such as locomotor activity, body temperature, O_2_ consumption (VO_2_), and respiratory quotient (RQ) showed significant diurnal rhythms in both conditions (Fig. 1C – F, Table S1). No marked differences in locomotor activity were seen between the groups (Fig. 1C, S1E). In contrast, in the T_3_ group, body temperature was elevated in the light (rest) phase (Fig. 1D, S1F) leading to a marked reduction in diurnal amplitude. Oxygen consumption in T_3_ was elevated throughout the day, but this effect was more pronounced during the dark phase (Fig. 1E, S1G) leading to an increase in diurnal amplitude. Linear regression of energy expenditure (EE) against body weight in CON and T_3_ mice (Tschöp et al. 2012) revealed no difference in slope, but a higher elevation/intercept was found in T_3_ mice (Fig. S1H). These data suggest that the higher EE of T_3_ mice is not only a consequence of increased body weight, but it also arises from a higher metabolic state. In T_3_ mice, RQ was slightly higher in the second half of the dark and the beginning of the light phase indicating higher carbohydrate utilization during this period (Fig. 1F, S1I). In summary, TH dependent changes in overall metabolic activity were observed resembling the human hyperthyroid condition, albeit with marked diurnal phase-specific effects.

These findings prompted us to evaluate to which extent T_3_ and T_4_ levels would be predictive for overall metabolic state (*TH state effects*) or, alternatively, for changes in metabolic activity across the day (*temporal TH effects*) by correlating hormone levels with metabolic parameters. When comparing daily averages to assess TH state effects, we found an association between T_3_ levels, body temperature and VO_2_ levels but not activity (Fig. 1G – I). Regarding temporal TH effects, we found that neither T_3_ nor T_4_ qualified as markers for diurnal variations in energy metabolism (Fig. 1J – O, Table S2). In summary, our data suggest that T_3_ levels are valid predictors of baseline metabolic state but fail to mirror diurnal changes in metabolic activity at, both, physiological and high-T_3_ states. T_4_ is an overall poor metabolic biomarker.

### Daytime-independent effects of TH on the liver transcriptome

To study the molecular pattern underlying the observed diurnal modulation of metabolic activity in T_3_-treated mice, we focused on the liver as a major metabolic tissue. We initially identified time-of-day independent transcriptional markers reflecting TH state in this tissue. Comparing the liver transcriptome across times of day and T_3_ treatment conditions, 2,343 differentially expressed probe sets (2,336 genes – DEGs) were identified (± 1.5-fold change; FDR < 0.1; Fig. 2A, Table S3). Of these DEGs, 1,391 and 945 genes were up-or downregulated, respectively, by elevated T_3_ (Fig. 2A, Table S3). Gene set enrichment analysis (GSEA) of upregulated DEGs yielded processes related to xenobiotic metabolism/oxidation-reduction, immune system, and cholesterol metabolism, amongst others. On the other hand, GSEA of downregulated DEGs yielded biological processes pertaining to fatty acid (FA) and carbohydrate metabolism, as well as cellular responses to insulin (Fig. 2B, Table S3). We identified 37 genes whose expression was robustly up-or down-regulated by T_3_ across all timepoints (Fig 2C, Table S4). Genes involved in xenobiotic transport/metabolism (*Abcc3, Abcg2, Ces4a, Ugt2b37, Papss2, Gstt1, Sult1d1, Cyp2d12, Ephx2*, and *Slc35e3*), lipid, fatty acid and steroids metabolism, (*Cyp39a1, Ephx2, Akr1c18, Acnat1, Cyp4a12a/b, Cyp2c44*), vitamin C transport (*Slc23a1*), and vitamin B_2_ (*Rfk*) and glutathione metabolism (*Glo1*) were identified. Additional genes involved in mitosis and replication were also identified (*Cep126, Mdm2, Trim24, and Mcm10*) (Fig. 2D, Table S4).

**Figure 2:**
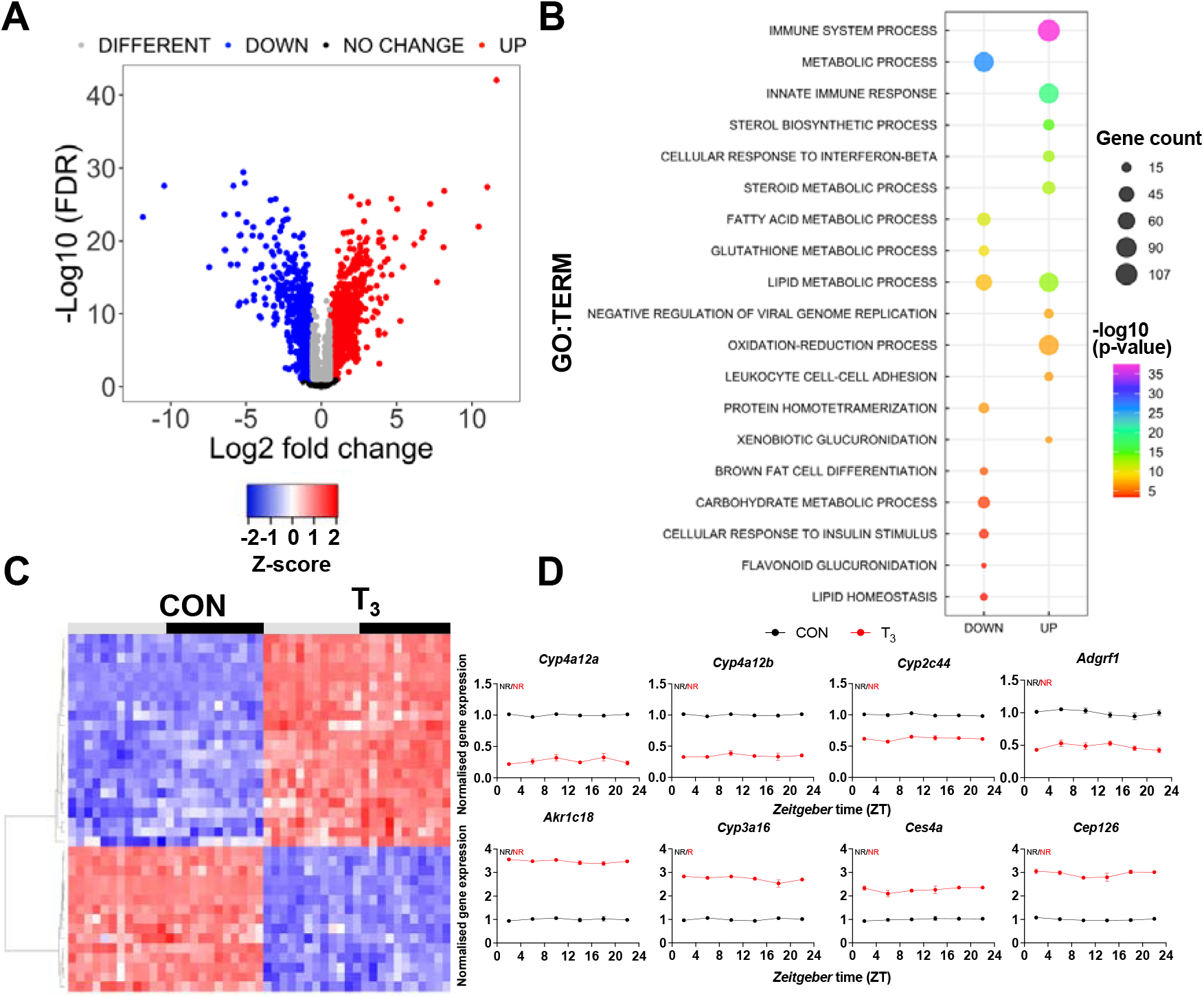
Identification of daytime-independent differentially expressed genes (DEGs) in liver of T_3_ mice. A) Global evaluation of liver transcriptomes revealed 2,336 DEGs of which 1,391 and 945 were considered as up-or downregulated, respectively, using a false discovery rate (FDR) < 0.1. Genes with an FDR < 0.1 were classified as different irrespectively of fold change values. B) Top-10 list of biological processes from gene set enrichment analyzes (GSEA) of up-and down-regulated DEGs are represented. Additional processes can be found in table S3. C) Heat map of liver DEGs showing significant T_3_-dependent regulation across all time points. Light and dark phases are shown as gray and black, respectively. D) Diurnal expression profiles of most robustly regulated DEGs. Gene expression of all both groups were normalized by CON mesor. Additional information is described in Table S4. None of these genes showed rhythmic regulation across the day (NR). n = 4 samples per group and timepoint, except for T_3_ group at ZT 22 (n =3).

We suggest that these transcripts could serve as robust daytime-independent biomarkers of TH state in liver.

### TH dependent regulation of liver diurnal transcriptional rhythms

We used the JTK cycle algorithm (Hughes et al. 2010) to describe the effects of TH state changes on 24-hour liver gene expression rhythms. We identified 3,354 and 2,592 probes – comprising 3,329 and 2,585 unique genes – as significantly rhythmic (p < 0.05) in CON or T_3_, respectively (Fig 3A, Table S5). Of these, 2,319 and 1,557 probes were classified as exclusively rhythmic in CON or T_3_, respectively. One thousand and thirty-five (1,035) probes (1,032 genes) were identified as rhythmic in both groups (Fig. 3A, Table S5), amongst these most core circadian clock genes (Table S5). Principal component analysis (PCA) showed a distinct pattern of organization across time between the groups for the shared genes (Fig. S2). We next assessed the distribution of phase and amplitude across 24 h between the groups. Rose plot analyzes revealed a similar distribution pattern of phase, but T_3_ mice showed a higher number of genes peaking in the light phase (ZT 7 – 9) and first half of dark phase (ZT 13 – 20) compared to CON (Fig. 3B). Cross-condition comparison of genes with robust rhythmicity revealed only a minor phase advance of around 1h in T_3_ (Fig. 3 C).

**Figure 3:**
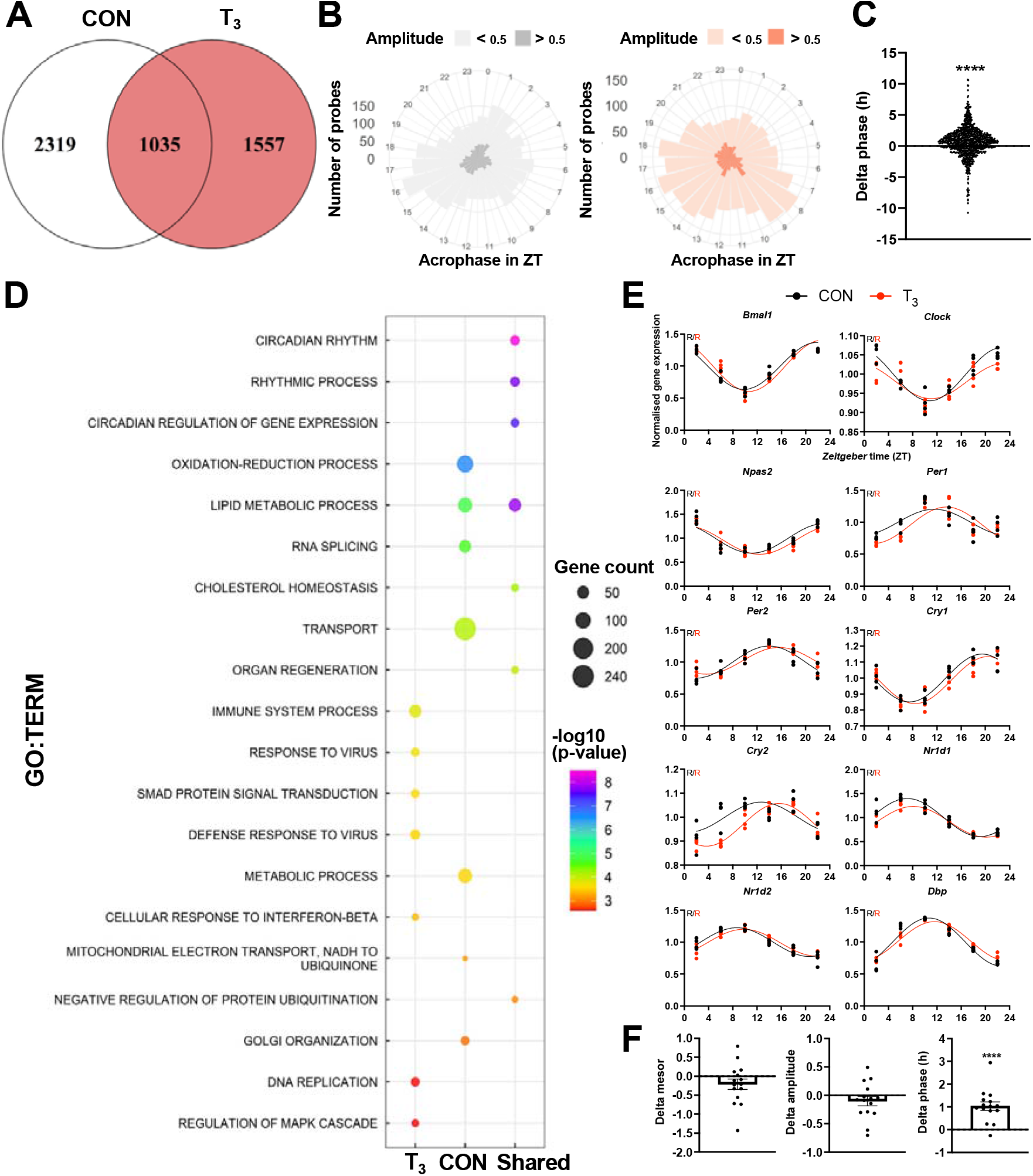
Diurnal evaluation of liver transcriptome of T_3_ mice. A) Rhythmic probes were identified using JTK cycle algorithm (Table S5). Venn diagram represents the distribution of rhythmic probes for each group. B) Roseplot of all rhythmic genes from CON (grey) and T_3_ (red) are represented by the acrophase and amplitude. Phase estimation was obtained from CircaSingle algorithm. C) Phase difference between shared rhythmic genes is shown. Each dot represents a single gene. One-sample *t* test against zero was performed and a significant interaction (mean 0.7781, p < 0.001) was found. D) Top 7 GSEA of exclusive genes from CON, T_3_, and shared are depicted. Additional processes are shown in Table S5. E) Sine curve was fitted for selected clock genes. Gene expression of all both groups were normalized by CON mesor. F) For mesor, amplitude, and phase delta assessment, CircaCompare algorithm was used. CON group was used as baseline. Additional genes (*Per3, Rorc, Tef, Hif1a*, and *Nfil3*) were used for these analyzes. 1-sample *t* test against zero value was used and only phase was different from zero (mean 1.036, p < 0.001). n = 4 samples per group and timepoint, except for T_3_ group at ZT 22 (n =3).

GSEA of rhythmic genes was performed to detect rhythmically regulated pathways under both TH conditions. In CON mice, transport, RNA splicing, lipid and glucose metabolism, and oxidation-reduction processes were overrepresented. In the high-T_3_ condition, several immune-related processes, fatty acid oxidation, and regulation of Mitogen-Activated Protein Kinase 1 (MAPK) signaling were found. Interestingly, robustly rhythmic genes were enriched for lipid and cholesterol metabolism and circadian related processes, suggesting that these processes are tightly coupled to circadian core clock regulation (Fig. 3D, Table S5). Individual inspection of clock genes revealed the absence of marked effects on mesor and amplitude, but a slight phase advance (Fig. 3E – F), which corroborates the phase advance effects seen at the rhythmic transcriptome level (Fig. 3C).

We next focused on the diurnal regulation of TH signaling by analyzing the expression of genes encoding for modulators of TH signaling, i.e., TH transporters, deiodinases, and TH receptors, and established TH target genes. We found that the TH transporter genes, *Slc16a2* (*Mct8*), *Slc7a8* (*Lat2*), and *Slc10a1* (*Ntcp*) lost rhythmicity in T_3_ mice compared to CON. Amongst the receptors, *Thra* was rhythmic, while *Thrb* was arrhythmic under both conditions. Of the deiodinases, only *Dio1* was robustly expressed under both conditions, but without variation across the day (Fig. 4A). Significant, but non-uniform changes in baseline expression levels were observed for *Slc16a10, Slc7a8, Dio1* (up in T_3_) and *Slco1a1, Thra*, and *Thrb* (down in T_3_; Fig. 4A). To analyze the effect of such changes on TH action, we studied diurnal regulation of established liver TH output genes. Reflecting elevated T_3_, all selected TH target genes showed increased expression across the day in T_3_ mice (Fig. 4B – C). No clear regulation was seen regarding amplitude or phase (Fig. 4 C).

**Figure 4:**
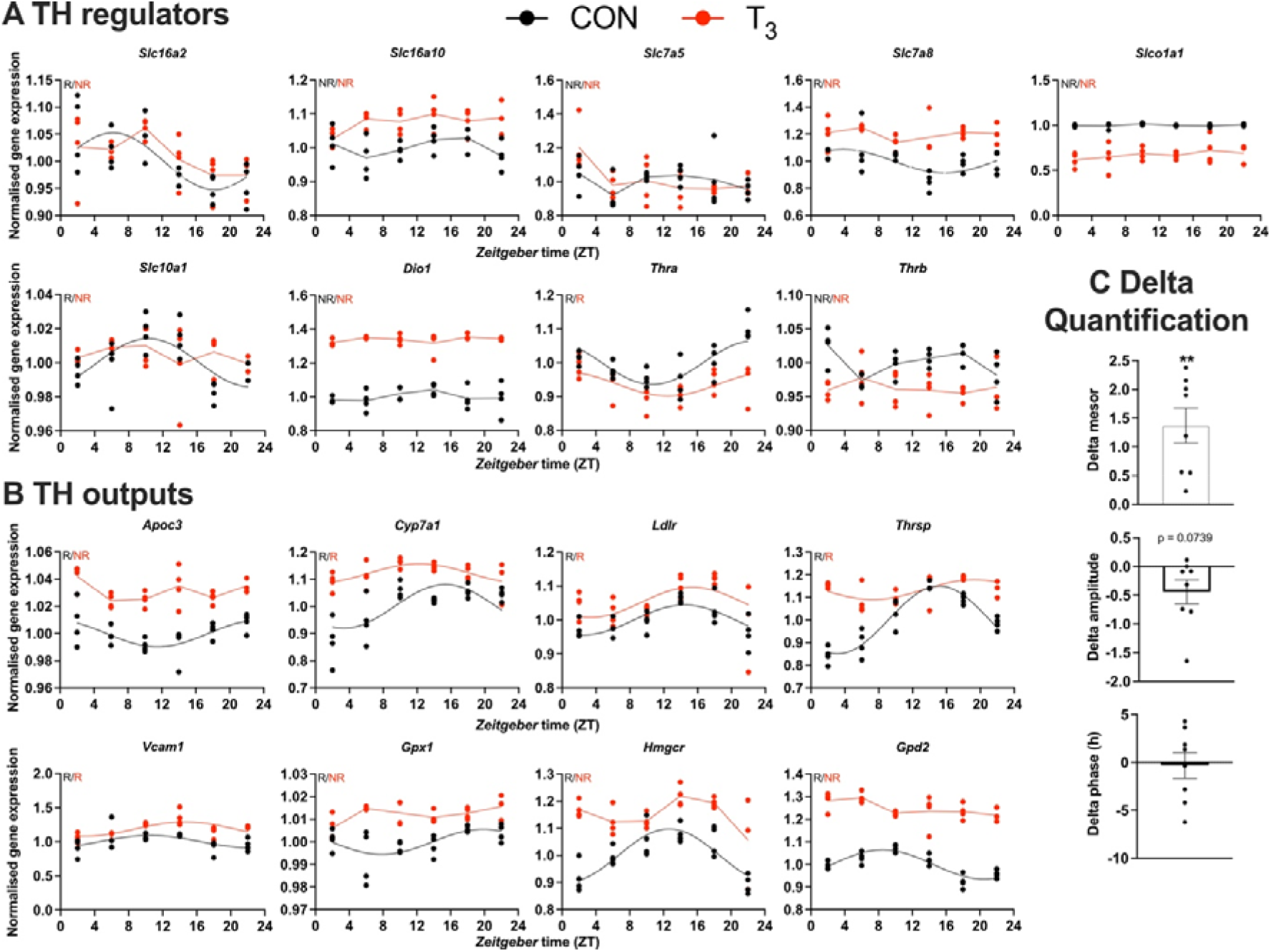
Gene expression evaluation of thyroid hormones (THs) regulators and metabolic outputs in T_3_ compared to CON. A and B) Genes involved in TH regulation, including transporters, *Dio1*, TH receptors, and well-known T_3_ outputs are presented. Presence (R) or absence of circadian rhythm (NR) detected by CircaCompare is depicted. Sine curve was fitted for rhythmic genes. Gene expression of all both groups were normalized by CON mesor. C) Evaluation of rhythmic parameters from genes described in B was performed by CircaCompare using CON group as baseline. 1-sample *t* test against zero value was used and only mesor was different from zero (mean 1.371, p < 0.01). n = 4 samples per group and timepoint, except for T_3_ group at ZT 22 (n =3).

In summary, we provide evidence that the molecular clock of the liver functions independent of TH state. At the same time, changes in diurnal expression patterns were found for fatty acid oxidation-and immune system-related genes in T_3_ mice. These changes were associated with marked gene expression profile alterations for TH signal regulators and outputs. Collectivity, these data indicate an adaptation of the diurnal liver transcriptome in response to changes in TH state in a largely tissue clock-independent manner.

### Quantitative characterization of TH dependent changes in liver diurnal transcriptome rhythms

To dissect TH state dependent rhythm alterations in the liver transcriptome, we employed CircaCompare (Parsons et al. 2020) to assess mesor and amplitude in genes that were rhythmic in at least one condition. For precise phase estimation, analyzes were performed only on robustly rhythmic genes. Of note, some differences in rhythm classification between JTK and CircaCompare were detected, which is expected due to the different statistical methods. Since we used CircaCompare’s rhythm parameter estimations for quantitative comparisons, gene rhythmicity cut-offs in the following analyzes were taken from this algorithm. Pairwise comparisons of rhythm parameters (*i*.*e*., mesor, amplitude, and phase) revealed predominant effects of TH state on mesor (2,519 probes / 2,504 genes) followed by alterations in amplitude (518 probes / 516 genes) and phase (491 probes/genes; Fig 5A, Table S6).

**Figure 5:**
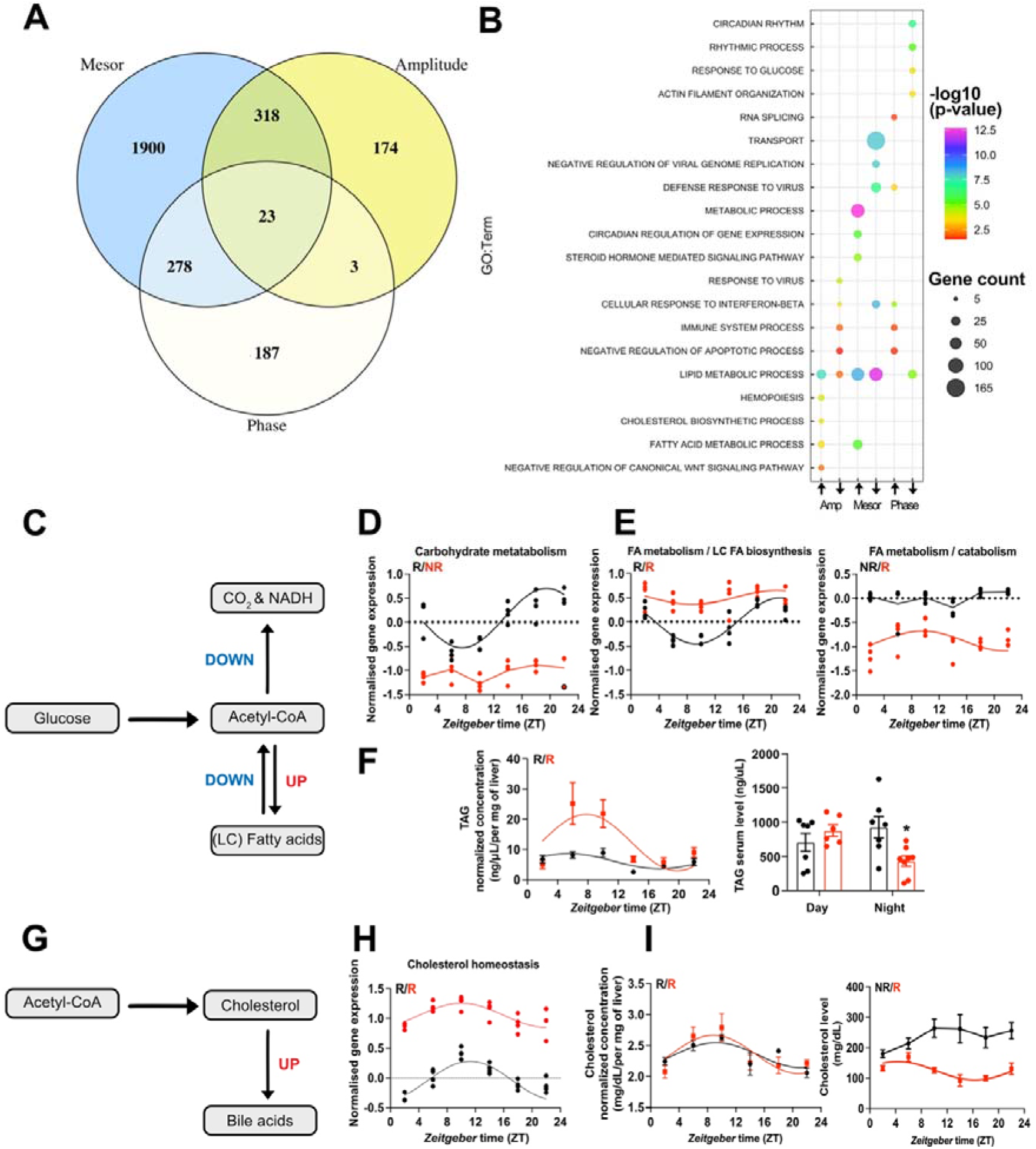
CircaCompare analyses of T_3_ (red) mice compared to CON (black). A) Venn diagram demonstrates the number of probes that displayed differences in each rhythmic parameter (mesor, amplitude, and phase). B) Top 5 enriched biological processes for each rhythmic parameter category. C) Summary of the CircaCompare analyzes regarding glucose and FA metabolism. D – E) Representation of glucose and fatty acid metabolism biological processes obtained from transcriptome data. F) Diurnal rhythm evaluation of liver TAG and day (ZT 2-6) vs night (ZT18-22) serum TAG levels comparisons. G) Summary of the CircaCompare analyzes regarding cholesterol metabolism. H) Representation of cholesterol homeostasis obtained from transcriptome data. I) Diurnal rhythm evaluation of liver and serum cholesterol. Gene expression from each biological process was averaged per ZT and plotted. The reader should refer to the text for detailed information regarding the changes found at the gene level of these processes. Sine curve was fitted for each rhythmic biological process. Individual gene expression pertaining to these processes is found in Fig. S3. n = 4 samples per group and timepoint, except for T_3_ group at ZT 22 (n =3).

We further differentiated CircaCompare outcomes into mesor or amplitude elevated (UP) or reduced (DOWN) and phase delayed or advanced for subsequent GSEA. In these analyzes, lipid metabolism was enriched in all categories, except for the phase advance group, which suggests a differential regulation of different gene sets related to lipid metabolism. GSEA of genes with reduced amplitude showed enrichment for fatty acid metabolism and cholesterol biosynthesis, whereas GSEA of elevated amplitude genes showed a strong enrichment for immune system-related genes. Interestingly, genes associated with circadian processes and response to glucose were enriched in the phase delay group (Fig. 5B, Table S6).

We extracted genes associated with glucose and fatty acid (FA) metabolic pathways from KEGG and assessed rhythmic parameter alterations according to CircaCompare (Fig. 5C – E, S3). Averaged and mesor-normalized gene expression data of each gene identified by GSEA were used to identify time-of-day dependent changes in biological processes.

Our data suggest a rhythmic pattern of glucose transport in CON mice roughly in phase with locomotor activity (Fig S3, Table S5; 1C). *Slc2a1* (*Glut1*) was rhythmic in both groups but showed a higher mesor in T_3_ mice (Fig. S3A; Table S5). Conversely, *Slc2a2* (*Glut2*), the main glucose transporter in the liver, was rhythmic in both groups, but it showed a reduced mesor in T_3_ mice. Other carbohydrate related transporters such as *Slc37a3* and *Slc35c1* gained rhythmicity and showed higher amplitude and/or mesor in T_3_ mice. Although GLUT1 role in liver is minor, increased GLUT1 signaling has been associated with liver cancer and in non-alcoholic steatosis (NASH) (Chadt and Al-Hasani 2020). CON mice, overall rhythmicity in carbohydrate metabolization transcripts, with acrophase in the dark phase, was identified whereas in T_3_ mice this process was arrhythmic due to a reduction in amplitude and mesor (Fig. 5D, S3A). Individual gene inspection showed that glucose kinase (*Gck*), an important gene that encodes a protein that phosphorylates glucose, thus allowing its internal storing and *Pgk1*, which encodes an enzyme responsible for the conversion of 1,3-diphosphoglycerate to 3-phosphoglycerate, showed reduction of amplitude in T_3_ mice. Loss of rhythmicity was found for *Pdk4*, a gene that encodes an important kinase that inhibits pyruvate dehydrogenase and for *Pdhb*, an important component of pyruvate dehydrogenase complex. Reduced inhibition of the pyruvate dehydrogenase complex is known to lead to less glucose utilization via tricarboxylic acid cycle and thus it favors ß-oxidation (Zhang, Hulver, et al. 2014).

Absence of rhythmicity and a higher mesor for the FA biosynthesis rate-limiting gene, *Fasn*, was found in T_3_ mice, despite this process was not enriched (Fig. S3B, Table S6). We identified two subsets of genes with a different regulation at mesor level in FA metabolism (Fig. 5E). Overall pathway analysis suggested reduced amplitudes associated with a higher mesor. Individual inspection revealed genes mainly related with unsaturated FA especially with biosynthesis (*Fads2*), and long chain FA elongation (*Elovl3, Acnat1-2*, and *Elovl6*), and oxidation (*Acox2*). *Fads2, Elovl2*, and *Elovl3* genes also showed a phase delay. Other subsets of genes showed reduced mesor without changes in amplitude, amongst these genes involved in FA biosynthesis (*Acsm3, Acsm5, Slc27a2*, and *Slc27a5*), ß-oxidation (*Acaa2, Hsd17b4, Crot, Acadl, Acadm, Hadh, Decr1, Cpt1a, Acsl1*, and *Hadhb*), glycerolipids biosynthesis (*Gpat4*), and FA elongation (*Hacd3*) (Fig. 5E, S3B, Table S6). To evaluate the metabolic consequences of T_3_ mediated diurnal rewiring of FA-related transcripts, we measured TAG levels in the liver across the day. TAG levels were rhythmic with an acrophase in the light phase in both groups. However, high T_3_ levels resulted in a marked increase in amplitude and mesor, thus arguing for a pronounced TAG biosynthesis in the light phase, followed by a stronger reduction in the dark phase, which points to higher TAG consumption. Interestingly, in serum, TAG levels were reduced only in the night phase, likely as a result of the higher energy demands of T_3_ mice (Fig 5F; 1E; Table S6). Taken altogether, our data suggest a preferential effect of T_3_ to increase FA biosynthesis and oxidation and a reduction in glucose metabolization as energy source in the liver.

A marked diurnal transcription rhythm was observed for cholesterol metabolism genes in CON mice (Fig. 5G – H). In T_3_ mice, cholesterol biosynthesis associated genes were enriched in the amplitude down group, thus suggesting a weakening of rhythmicity. Within this line, the rate-limiting enzyme encoding gene, *Hmgcr*, showed loss of rhythmicity with reduced amplitude and increased mesor in T_3_ mice (Fig S3C; Table 6). Interestingly, upon evaluation of liver cholesterol levels no significant difference was observed, although in both groups, cholesterol levels were rhythmic and with an acrophase in the rest phase. In serum, only in T_3_ mice, cholesterol levels were rhythmic but showed a marked mesor reduction compared to CON, especially in the dark phase (Fig 5I; Table 6). Rhythmic genes with a marked higher mesor involved in cholesterol uptake (*Ldlr, Lrp5*, and *Nr1h2*) and secretion (*Abcg5*/8 and *Cyp7a1*) in bile acids (Fig 5G; S3C; Table 6) were detected in line with T_3_ mediated increased bile acid production (Gebhard and Prigge 1992; Bonde et al. 2012). Taken altogether, our data suggest T_3_ mediated time-restricted reduction of cholesterol serum levels in favor of increased cholesterol metabolization.

## DISCUSSION

In this study, we analyzed the effects of high T_3_ state in the mouse liver. Our data argue that T_3_ is a marker for time-independent metabolic output which is subject to distinct temporal (i.e., diurnal) modulation. At the transcriptome level, T_3_ induction led to metabolic pathway rewiring associated with only a minor impact on the circadian clock machinery of the liver.

Upon analyzing the diurnal metabolic effects of T_3_, we identified a reduction of core body temperature amplitude due to an elevation in the light phase. Conversely, T_3_ mice showed a higher O_2_ consumption amplitude due to increased respiratory activity in the dark phase. Day versus night analyzes confirmed that during the light phase, T_3_ mice have increase metabolic output, which become higher during the dark phase. The absence of an effect in locomotor activity between the groups, reinforces the fact of T_3_ as strong activator of energy metabolism in our study, which is support by experimental data (Lanni et al. 2005; Cioffi et al. 2010; Mullur et al. 2014; Jonas et al. 2015). Thus, one could suggest that several adaptive mechanisms must happen to increase basal metabolic rate. In this line, increased energy output shown by T_3_ mice seems to relay on a slightly increased glucose (higher RQ quotient) consumption both at light and dark phases. In the liver, our transcriptome analyzes revealed important changes in gene expression reflecting increased metabolic output, which will be discussed below.

Although daytime specific effects in metabolic outputs were observed, no clear correlation between TH levels and metabolic outputs was found, thus ruling out that T_3_ or T_4_ are useful *temporal* markers for metabolic output. On the other hand, as a *state* marker, i.e., when seen on a longer perspective, T_3_ served as a robust predictor of metabolic output. For T_4_, a lack of temporal correlation is easily explained by the absence of diurnal rhythmicity in both normal-and high-T_3_ conditions. Conversely, T_3_ levels were rhythmic in only CON mice, and thus, the lack of T_3_ correlational effect may reside in absence of rhythmicity in the T_3_ group. Previous studies have suggested that serum T_3_ shows lack of rhythmicity, or if it is present, displays rhythms of small amplitude in humans and/or mice (Weeke and Laurberg 1980; Russell et al. 2008; Philippe and Dibner 2015). In our experimental conditions, CON mice displayed a stable circadian rhythm of T_3_, albeit with a low amplitude.

Nonetheless, different set of genes were differentially expressed at different times of the day, thus suggesting time-dependent effects of T_3_ in the liver. This is suggestive of additional underlying mechanisms that are not dependent on the oscillatory T_3_ serum levels. We hypothesized that the liver could display increased sensitivity to T_3_ effects likely via rhythmicity in TH transporters, *Dio1*, and TH receptors expression and/or activity. To illustrate this concept, our transcriptome analyzes showed that the liver diurnal transcriptome has 2,336 robustly regulated genes (ca.10% of the transcriptome). Previous studies from the early 2000s using microarrays identified about 2-5 % as T_3_ responsive genes (Feng et al. 2000; Flores-Morales et al. 2002). Experimental differences such as different T_3_ levels associated with differences in statistical and significance threshold levels contribute to the differences found between our data and the previous studies. Enrichment analyzes showed that elevated levels of T_3_ were associated with oxidation-reduction and immune system related genes whereas a negative association was found for glucose and FA metabolism.

Focusing on comprehending time of day dependent effects in the liver, we focused on the differently expressed genes per timepoint. We identified several hundreds of DEGs across time in T_3_ mice, thus arguing for a time-dependent effect of T_3_ in the liver. *Dio1* expression is classically associated with liver thyroid state (Zavacki et al. 2005). In our dataset, *Dio1* was differently expressed in all ZTs, except for ZT 22, an effect caused by increased variation in the CON group. Remarkably, 37 genes were identified as time-independent DEGs, i.e., displayed stable T_3_ state dependent expression across all time points, of which were 22 up-and 15 downregulated in T_3_ mice. These genes participate in several biological processes such as xenobiotic transport/metabolism, lipid, fatty acid metabolism, and amongst others. From a translational view, we suggest that these genes could be used to evaluate the thyroid state of the liver at any given time in experimental studies. Moreover, these genes could be used to create a signature of thyroid state in the liver in different conditions and diseases.

While tonic transcriptional targets of T_3_ have been described in tissues such as the liver, at the same time, robust diurnal regulation of modulators of thyroid hormone action such as TH transporters, deiodinases, and TH receptors can be observed from high-resolution circadian studies ((Zhang, Lahens, et al. 2014); http://circadiomics.igb.uci.edu). This prompted us to study how T_3_ may affect the transcriptional outputs across the day using established circadian biology methods. Circadian evaluation of CON and T_3_ livers revealed 10 – 15 % of the liver transcriptome as rhythmic under both experimental conditions, which is in line with previous experiments (Zhang, Lahens, et al. 2014; Greco et al. 2021). 1,032 genes (ca. 5 % of the liver transcriptome) were robustly rhythmic under both T_3_ conditions. Overall, the elevation of T_3_ had a slight phase delaying effect on these rhythmic genes, which is similar to the effects found in core circadian clock genes.

mRNA expression of TH transporter genes, *Slc16a2* (*Mct8*), *Slc7a8* (*Lat2*), and *Slc10a1* (*Ntcp*), showed a loss of rhythmicity while no gain of rhythmicity was found for T_3_ mice. Such loss of rhythmicity in TH transporters could represent a compensatory mechanism to the higher T_3_ levels found across the day. The transcriptional response in TH regulators suggests a desensitization mechanism in the liver of T_3_ mice with a downregulation of TH receptors but increased baseline expression of *Dio1, Slc16a10*, and *Slc7a8*. Collectively, these data suggest a compensatory mechanism of decreased signal responses, elevated transport and metabolization of T_3_ under high-T_3_ conditions at the mRNA level. However, one must consider the potential diurnal regulation of TH receptor protein levels as well as DIO1 and transporter activity to fully confirm this putative compensatory mechanism.

Regarding diurnal changes, we observed a strong effect of T_3_ on mesor, followed by changes in amplitude and phase. Interestingly, while circadian parameter analysis revealed a strong effect of T_3_ on liver transcriptome rhythms, this was mostly without affecting the molecular clock machinery itself. Therefore, T_3_ effects in the liver seem to act downstream of the molecular clock through a still elusive mechanism.

Considering the broad range of changes found in our study, we focused our efforts on comprehending T_3_ effects on metabolic pathways. Our data reveal a strong T_3_ mediated diurnal regulation of energy metabolism, mainly related to glucose and FAs, on the mRNA level. Transcripts associated with both processes lost their rhythmicity under high-T_3_ conditions, thus becoming constant across the day. Interestingly, we found evidence that T_3_ leads to a shift towards FA ß-oxidation over glucose utilization in the liver. T_3_ effects in FA biosynthesis showed a preferential effect on the synthesis and oxidation of long chain FA on the mRNA level. Confirming our predictions, livers from T_3_ mice had higher levels of TAG during the light phase compared to CON, thus suggesting a higher TAG synthesis during the rest phase. However, during the dark phase a marked reduction in TAG serum and liver levels were observed, which suggests an important role of FA ß-oxidation as energy source to meet the higher energetic demands imposed by T_3_. Indeed, such changes can be associated with higher energetic demands (higher VO_2_) both during the light and dark phase in T_3_ mice. It is a known fact that T_3_ increases TAG synthesis in the liver (Sinha et al. 2018), but our data provide an interesting time of day dependency in T_3_ effects. Interestingly, no marked alteration in protein catabolism was found, thus suggesting a preferential effects of T_3_ for glucose and FA related energy sources, at least in liver (Mullur et al. 2014).

Our bioinformatic analyzes predicted a higher pool of acetyl-CoA in liver of T_3_ mice as consequence of higher FA ß-oxidation, which we hypothesized being associated with a putative increased cholesterol biosynthesis. However, no differences were observed in liver cholesterol between the groups, but a marked reduction in serum cholesterol levels was identified in T_3_ mice. In face of no differences in cholesterol levels in liver, but associated with a marked reduction in serum cholesterol, we suggest a cholesterol higher uptake and conversion into bile acid. Indeed, such mechanism is supported by our transcriptomic data as well as the literature as T_3_ is known to increase cholesterol secretion via bile acids or non-esterified cholesterol in the feces (Mullur et al. 2014; Sinha et al. 2018). Such marked diurnal alterations in the liver transcriptome, especially with regards to metabolic pathways, led us to speculate on the overall consequences of high T_3_ on organismal rhythms. Loss or weakening of rhythmicity in relevant metabolic processes in other organs, such as the pancreas, white and brown adipose tissue, and other organs, may also take place in the high-T_3_ condition, which could explain the higher energetic demands induced by elevated T_3_ levels. It is still elusive how T_3_ affects other metabolic and non-metabolic organs in a circadian way. Such knowledge will proof useful in design therapeutic strategies for TH-related diseases such as hepatic steatosis (Marjot et al. 2022).

Considering the effects seen in the liver circadian transcriptome, associated with the metabolic data provided, we suggest that T_3_ may act as a rewiring factor of metabolic rhythms. In this sense, T_3_ leads to reduction of rhythmicity of major metabolic pathways to sustain higher energy demands across the day. Such pronounced effects are not reflected in marked alterations in the liver clock. From a chronobiological perspective, T_3_ may be considered a disruptor that uncouples the circadian clock from its outputs, thus promoting a state of chronodisruption (Potter et al. 2016; de Assis and Oster 2021). This duality of T_3_ effects warrants further investigation.

An exciting concept that arises from our data is the concept of chrono-modulated regimes for thyroid-related diseases such as hypo-and hyperthyroidism. We suggest evidence that the liver and presumably other organs may show temporal windows in which treatment can be more effective. Based on our diurnal transcriptome data, no optimal time could be suggested due to the lack of rhythmicity for *Dio1, Thrb*, and other TH regulators genes. Nonetheless, time-dependent effects in other genes and/or biological processes were identified and could be explored for chronotherapeutic drug intervention. Taken altogether, our study shows that T_3_ displays time of day dependent effects in metabolism output and liver transcriptome, despite the presence of a strong T_3_ diurnal rhythm. With regards to metabolism, T_3_ acts as a *state* marker, but fails to reflect temporal regulation of metabolic output. Metabolic changes induced by T_3_ resulted in a higher overall activation and loss of rhythmicity of genes involved in glucose and FA metabolism, concomitant with higher metabolic turnover, and independent of the liver circadian clock. Collectively, our data suggest a novel layer of diurnal regulation of liver metabolism that can bear fruits for future treatments of thyroid related diseases.

## MATERIAL AND METHODS

### Mouse model and experimental conditions

Two to three months-old male C57BL/6J mice (Janvier Labs, Germany) were housed in groups of three under a 12-hour light, 12-hour dark (LD, ∼300 lux) cycle at 22 ± 2 °C and a relative humidity of 60 ± 5 % with *ad-libitum* access to food and water. To render mice hyperthyroid (i.e., high T_3_ levels) the animals received one week of 0.01 % BSA (Sigma-Aldrich, St. Louis, USA, A7906-50G) in their drinking water, followed by two weeks with water supplemented with T_3_ (0.5 mg/L, Sigma-Aldrich T6397, in 0.01 % of BSA). Control animals received only 0.01 % BSA in the drinking water over the whole treatment period. During the treatment period mice were monitored for body weight and rectal temperature (BAT-12, Physitemp, Clifton, USA) individually and food and water intake per cage. All *in vivo* experiments were ethically approved by the Animal Health and Care Committee of the Government of Schleswig-Holstein and were performed according to international guidelines on the ethical use of animals. Sample size was calculated using G-power software (version 3.1) and are shown as biological replicates in all graphs. Experiments were performed at three to four times. Euthanasia was carried out using cervical dislocation and tissues were collected every 4 h. Night experiments were carried out under dim red light. Tissues were immediately placed on dry ice and stored at -80 °C until further processing. Blood samples were collected from the trunk, and clotting was allowed for 20 min at room temperature. Serum was obtained after centrifugation at 2,500 rpm, 30 min, 4 °C and samples stored at -20 °C.

### Total T_3_ and T_4_ evaluation

Serum quantification of T_3_ and T_4_ was performed using commercially available kits (NovaTec, Leinfelden-Echterdingen, DNOV053, Germany for T_3_ and DRG Diagnostics, Marburg, EIA-1781, Germany for T_4_) following the manufactures’ instructions.

### Serum and tissue triacylglycerides (TAG) and cholesterol evaluation

TAG and cholesterol evaluation of tissue and serum were processed according to the manufactures’ instructions (Sigma-Aldrich, MAK266 for TAG and Cell Biolabs, San Diego, USA, STA 384 for Cholesterol).

### Telemetry and metabolic evaluation

Core body temperature and locomotor activity were monitored in a subset of single-housed animals using wireless transponders (E-mitters, Starr Life Sciences, Oakmont, USA). Probes were transplanted into the abdominal cavity of mice 7 days before starting the drinking water treatment. During the treatment period mice were recorded once per week for at least two consecutive days. Recordings were registered in 1-min intervals using the Vital View software (Starr Life Sciences). Temperature and activity data were averaged over two consecutive days (treatment days: 19/20) and plotted in 60-min bins.

An open-circuit indirect calorimetry system (TSE PhenoMaster, TSE Systems, TSE Systems, USA) was used to determine respiratory quotient (RQ = carbon dioxide produced / oxygen consumed) and energy expenditure in a subset of single-housed mice during drinking water treatment. Mice were acclimatized to the system for one week prior to starting the measurement. Monitoring of oxygen consumption, water intake as well as activity took place simultaneously in 20-min bins. VO_2_ and RQ profiles were averaged over two consecutive days (treatment days: 19/20) and plotted in 60-min bins. Energy expenditure was estimated by determining the caloric equivalent according to Heldmaier (Heldmaier 1975): heat production (mW) = (4.44 + 1.43 * RQ) * VO_2_ (ml O_2_/h). A linear regression between EE and body weight was performed to rule out a possible confounding factor of body weight (Tschöp et al. 2012).

### Microarray analysis

Total RNA was extracted using TRIzol (Thermofisher, Waltham, USA) and the Direct-zol RNA Miniprep kit (Zymo Research, Irvine, USA) according to the manufacturer’s instructions. Genome-wide expression analyses was performed using Clariom S arrays (Thermo Fisher Scientific) using 100 ng RNA of each sample according to the manufacturer’s recommendations (WT Plus Kit, Thermo Fisher Scientific). Data was analyzed using Transcriptome Analyses Console (Thermo Fisher Scientific, version 4.0) and expressed in log_2_ values.

### Differentially expressed gene (DEG) analysis

To identify global DEGs, all temporal data from each group was considered and analyzed by *Student’s t* test and corrected for false discovery rates (FDR < 0.1). Up-or downregulated DEGs were considered when a threshold of 1.5-fold (0.58 in log_2_ values) regulation was met. As multiple probes can target a single gene, we curated the data to remove ambiguous genes. To identify DEGs at specific time points (ZTs – Zeitgeber time; ZT0 = “lights on”), the procedure described above for each ZT was performed separately. Time-independent DEGs were identified by finding consistent gene expression pattern across all ZTs.

### Rhythm analysis

To identify probes that showed diurnal (i.e., 24-hour) oscillations, we employed the non-parametric JTK_CYCLE algorithm (Hughes et al. 2010) in the Metacycle package (Wu et al. 2016) with a set period of 24 h and an adjusted p-value (ADJ.P) cut-off of 0.05. For visualization, data were plotted in Prism 9.0 (GraphPad, USA) and a sine wave was fit with a period set at 24 h. Rhythmic gene detection by JTK_CYLCE was evaluated by CircaSingle, a non-linear cosinor regression included in the CircaCompare algorithm (Parsons et al. 2020), largely (ca. 99 %) confirming the results from JTK_CYCLE. Phase and amplitude parameter estimates from CircaSingle were used for rose plot visualizations. To directly compare rhythm parameters (mesor and amplitude) in gene expression profiles between T_3_ and CON, CircaCompare fits were used irrespective of rhythmicity thresholds. Phase comparisons were only performed when a gene was considered as rhythmic in both conditions (p < 0.05).

### Gene set enrichment analysis (GSEA)

Functional enrichment analysis of DEGs was performed using the Gene Ontology (GO) annotations for Biological Processes on the Database for Annotation, Visualization, and Integrated Discovery software (DAVID 6.8 (Huang et al. 2009). Processes were considered significant for a biological process containing at least 5 genes (gene count) and a p-value < 0.05. To remove the redundancy of GSEA, we applied the REVIGO algorithm (Supek et al. 2011) using default conditions and a reduction of 0.5. Enrichment analyzes from genes sets containing less than 100 genes, biological processes containing at least 2 gene were included. Overall gene expression evaluation of a given biological process was performed by normalizing each timepoint of CON and T_3_ by CON mesor. A sine curve was plot and used for representation of significantly rhythmic profiles.

### Principle component analysis (PCA) plots

For PCA analyzes, each timepoint was averaged to a single replicate and analyzes were performed using the factoextra package in R and Hartigan-Wong, Lloyd, and Forgy MacQueen algorithms (version 1.0.7).

### Data handling and statical analysis of non-bioinformatic related experiments

Samples were only excluded upon technical failure. For temporal correlation analyzes, normalized values were obtained by dividing each value by the daily group average. Normalized values were correlated with normalized T_3_ and T_4_ levels using Spearman’s correlation. Correlation analyzes were performed between different groups of animals that underwent the same treatment. Analyzes were done in Prism 9.0 (GraphPad) and a p-value of 0.05 was used to reject the null hypothesis. Data from ZT0-12 were considered as light phase and from ZT 12 to 24 as dark phase. Data were either averaged or summed as indicated. Temporal data between groups were analyzed by two-way ANOVA followed by Bonferroni post-test. Single timepoint data were evaluated by unpaired *Student’s* t test with Welch correction or Mann-Whitney test for parametric or non-parametric samples, respectively.

### Data handling and statical analysis of bioinformatic experiments

Statistical analyses were conducted using R 4.0.3 (R Foundation for Statistical Computing, Austria) or in Prism 9.0 (GraphPad). Rhythmicity was calculated using the JTK_CYLCE algorithm in meta2d, a function of the MetaCycle R package v.1.2.0 (Wu et al. 2016). Rhythmic features were calculated and compared among multiple groups using the CircaCompare R package v.0.1.1 (Parsons et al. 2020). Data visualization was performed using the ggplot2 R package v.3.3.5, eulerr R package v.6.1.1, and Prism 9.0 (GraphPad). Heatmaps were created using the Heatmapper tool (http://www.heatmapper.ca).

### Data availability

All experimental data are deposited in the <u>Figshare</u> depository. Microarray data was deposited in the Gene Expression Omnibus (GEO) database (<u>GSE199998</u>). Upon publication all datasets will be publicly available.

## ACKNOWLEDGEMENTS AND FUNDING

This work was supported by grants of the German Research Foundation (DFG) to HO 353-10/1, GRK-1957, and CRC-296 “LocoTact” (TP13 and TP14). JTH is a fellow of the São Paulo Research Foundation (FAPESP - 04524-8/2020). We thank Lucas Moreira Ribeiro from the Federal University of Ouro Preto (UFOP, Brazil) for technical assistance in the bioinformatics analyzes.

## AUTHOR CONTRIBUTIONS

LH, JM, and HO conceptualization. LVMA data curation. LH and LVMA formal analysis and investigation. LH, LVMA, JTL, RP, MK, IC, IN, and OR, methodology. HO funding acquisition, project administration, and supervision. LVMA and HO writing - original draft. All authors: text review & editing.

---

**Figure S1:**
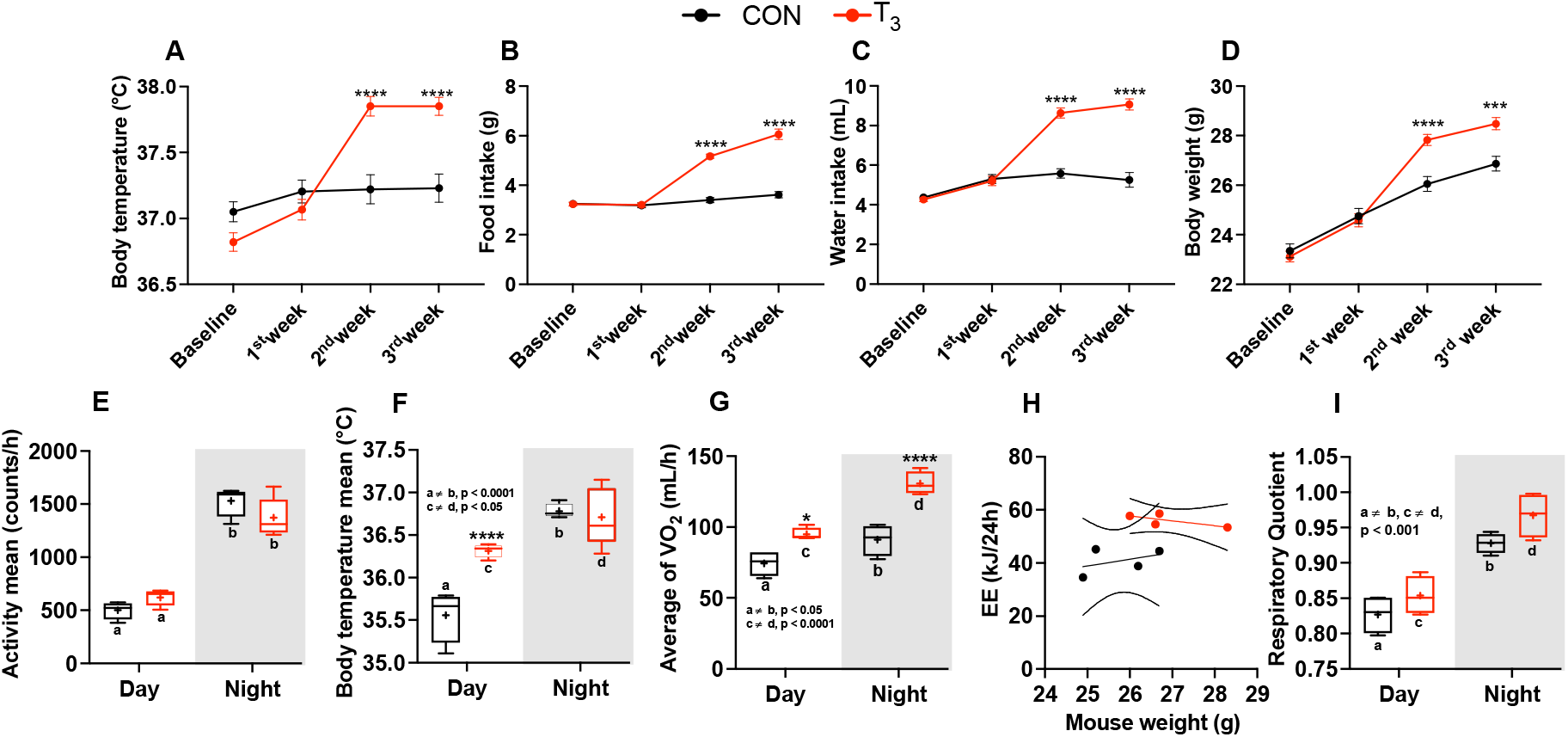
Metabolic evaluation of CON and T_3_ mice. A – D) Assessment of body temperature, food and water intake (per cage, n = 8), and body weight. E – I) Metabolic parameters (described in the y-axis) were obtained from the 3_rd_ week of experiment (days 19/20). Day and night data were obtained by averaging values from ZT 0 to 12 (day) and from ZT 12 to 24 (night) and plot accordingly. Letters represent a difference between the same group in day versus night comparisons. Asterisks represent significant differences between CON and T_3_ mice. In H, 95% confidence interval are shown. Comparison of the slope and elevations/intercept between the groups were performed: p = 0.30 and 0.01, respectively. Data are shown either as mean ± SEM or by boxplot. N = 24 for A and D. E – I) n = 4 – 5 per group.

**Figure S2:**
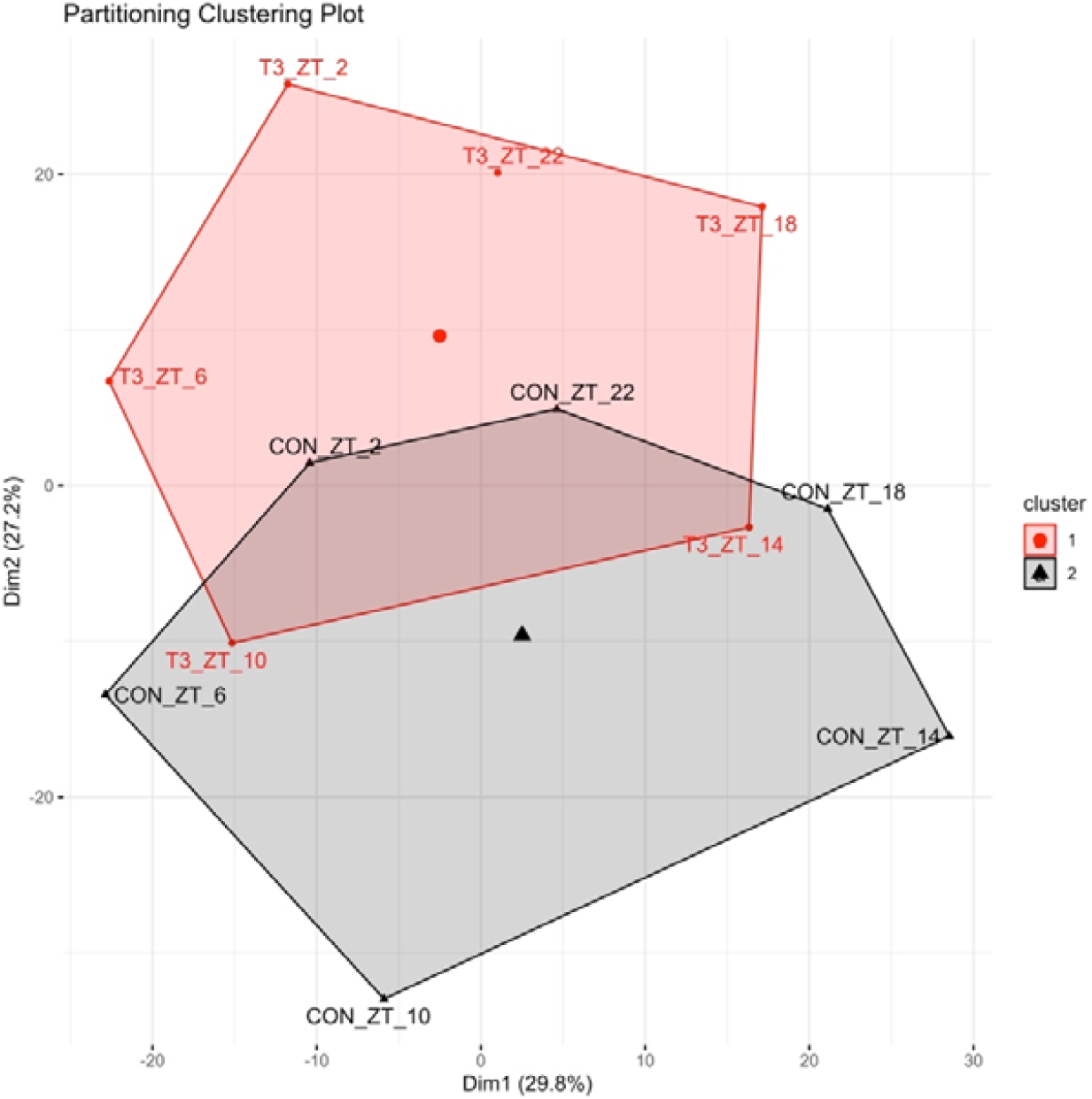
PCA plots of shared rhythmic genes. Each timepoint was averaged into a single replicate and PCA analyzes were performed using the factoextra package in R and Hartigan-Wong, Lloyd, and Forgy MacQueen algorithms.

**Figure S3.**
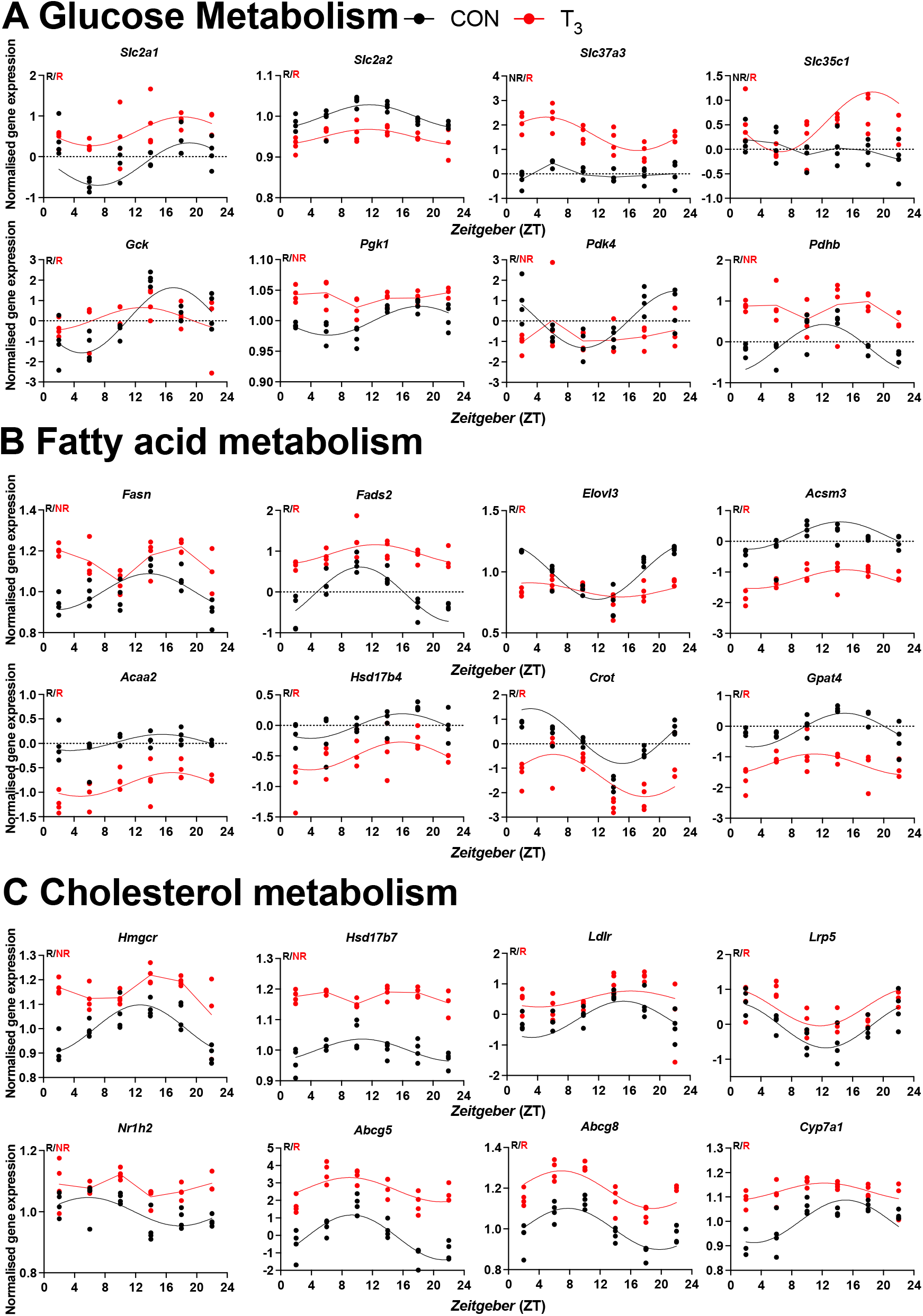
Expression profile of selected genes pertaining to biological processes identified in CircaCompare. Diurnal profile of genes from glucose (A), fatty acid (B) and cholesterol metabolism (C). Diurnal overall gene expression was normalized by CON mesor and plotted. Sine curve was fitted for rhythmic genes (R). Absence of rhythmic is represented by connected lines and NR symbol. n = 4 samples per group and timepoint, except for T_3_ group at ZT 22 (n =3). CircaCompare data is provided in Table S6.

